# Mixing the Message: Do Dung Beetles (Coleoptera: Scarabaeidae) Affect Dung-Generated Greenhouse Gas Emissions?

**DOI:** 10.1101/2020.11.19.389601

**Authors:** Fallon Fowler, Christopher J. Gillespie, Steve Denning, Shuijin Hu, Wes Watson

## Abstract

By mixing and potentially aerating dung, dung beetles may affect the microbes producing the greenhouse gases (GHGs): carbon dioxide (CO_2_), methane (CH_4_), and nitrous oxide (N_2_O). Here, their sum-total global warming effect is described as the carbon dioxide equivalent (CO_2_e). Our literature analysis of reported GHG emissions and statistics suggests that most dung beetles do not, however, reduce CO_2_e even if they do affect individual GHGs. Here, we compare the GHG signature of homogenized (“premixed”) and unhomogenized (“unmixed”) dung with and without dung beetles to test whether mixing and burial influence GHGs. Mixing by hand or by dung beetles did not reduce any GHG – in fact, tunneling dung beetles increased N_2_O medians by ≥1.8x compared with dung-only. This suggests that either: 1) dung beetles do not meaningfully mitigate GHGs as a whole; 2) dung beetle burial activity affects GHGs more than mixing alone; or 3) greater dung beetle abundance and activity is required to produce an effect.

## Introduction

Dung beetles (Coleoptera: Scarabaeidae) provide beneficial ecosystem services including improved nutrient recycling, competitive exclusion of pests and parasites, reduced animal disease incidence, and improved soil percolation and plant growth (Nichols et al. 2008). These arthropods also mitigate damaging ecological impacts such as water and soil pollution, and increased GHG production such as found in animal agriculture. These impacts are the result of excessive land-use, heavy reliance on fossil fuels, and concentrated animal waste that destroy and damage wildlife habitats (Steinfeld et al. 2006). In fact, animal agriculture produces 37% and 65% of global anthropogenic CH_4_ and N_2_O emissions (Steinfeld et al. 2006), with manure management accounting for 4.3% (not including deposited dung emissions) and ∼17% of livestock-produced CH_4_ and N_2_O, respectively (Gerber et al. 2013). Thriving dung beetle communities combat and sustainably restore dung-polluted habitats (Doube 2018), and so are ideal animals with which to study the GHG-resource recycling connection (Sylvia et al. 2005).

Recently, researchers investigated how dung beetles affect individual GHGs (Penttilä et al. 2013, Iwasa et al. 2015, Hammer et al. 2016a, Slade et al. 2016a, Piccini et al. 2017, Evans et al. 2019). Very often they reported net beneficial dung beetle effects (Iwasa et al. 2015, Hammer et al. 2016a, Slade et al. 2016a, Piccini et al. 2017) despite not reducing the total CO_2_e calculated from CH_4_, CO_2_, and N_2_O (Table 1).

**Table 1.**
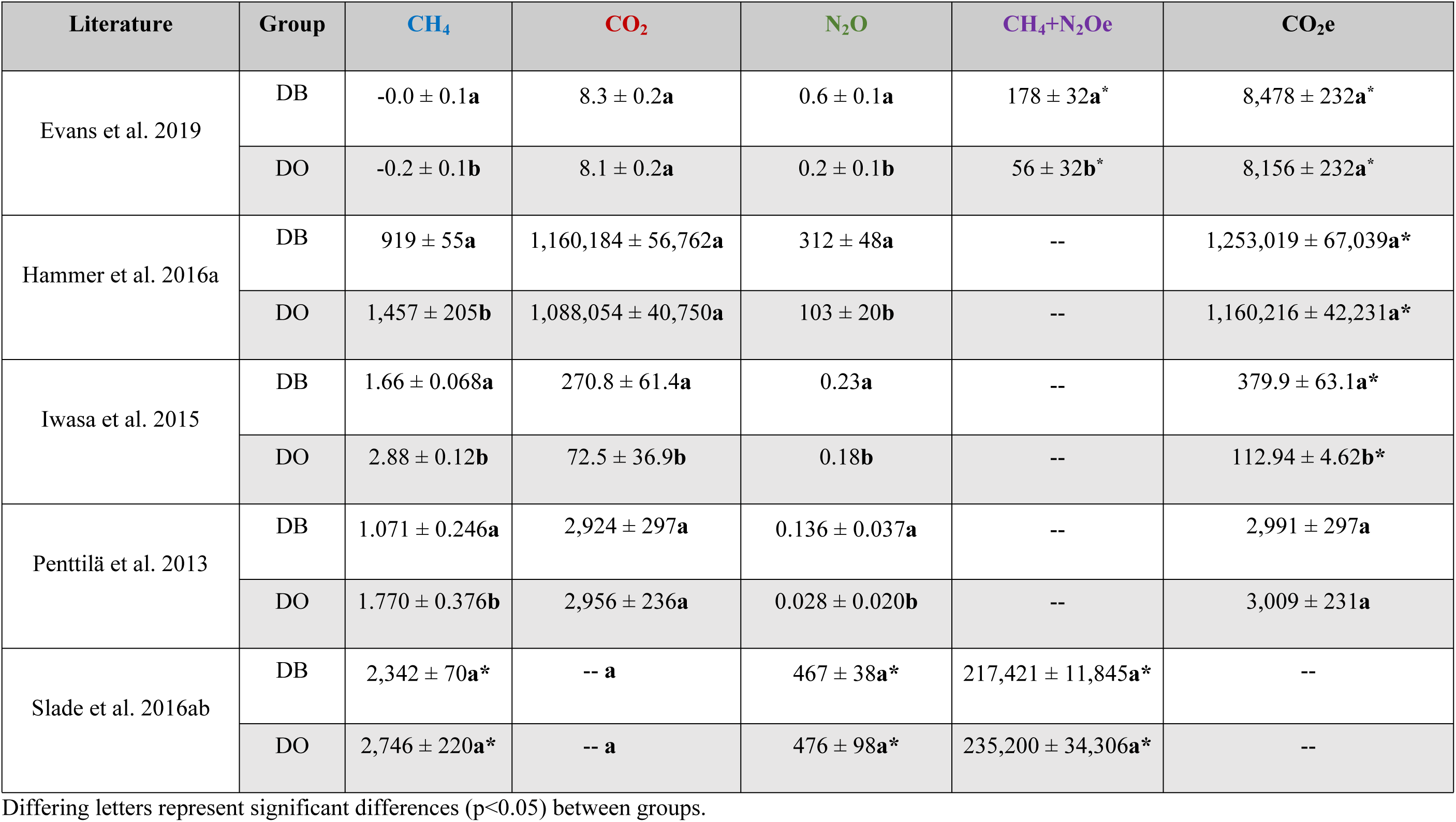
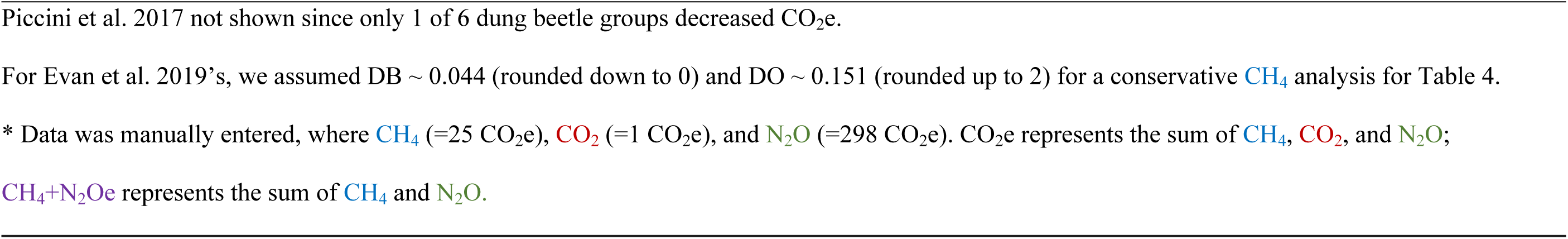
A summary of the exact GHG fluxes between dung beetles (DB) and dung-only (DO) from data and analyses of past authors.

| Literature | Group | CH<sub>4</sub> | CO<sub>2</sub> | N<sub>2</sub>O | CH<sub>4</sub>+N<sub>2</sub>Oe | CO<sub>2</sub>e |
| --- | --- | --- | --- | --- | --- | --- |
| Evans et al. 2019 | DB | -0.0 ± 0.1**a** | 8.3 ± 0.2**a** | 0.6 ± 0.1**a** | 178 ± 32 **a** * | 8,478 ± 232 **a** * |
|  | DO | -0.2 ± 0.1**b** | 8.1 ± 0.2**a** | 0.2 ± 0.1**b** | 56 ± 32 **b** * | 8,156 ± 232 **a** * |
| Hammer et al. 2016a | DB | 919 ± 55**a** | 1,160,184 ± 56,762**a** | 312 ± 48**a** | -- | 1,253,019 ± 67,039 **a** * |
|  | DO | 1,457 ± 205**b** | 1,088,054 ± 40,750**a** | 103 ± 20**b** | -- | 1,160,216 ± 42,231 **a** * |
| Iwasa et al. 2015 | DB | 1.66 ± 0.068**a** | 270.8 ± 61.4**a** | 0.23**a** | -- | 379.9 ± 63.1 **a** * |
|  | DO | 2.88 ± 0.12**b** | 72.5 ± 36.9**b** | 0.18**b** | -- | 112.94 ± 4.62 **b** * |
| Penttilä et al. 2013 | DB | 1.071 ± 0.246**a** | 2,924 ± 297**a** | 0.136 ± 0.037**a** | -- | 2,991 ± 297**a** |
|  | DO | 1.770 ± 0.376**b** | 2,956 ± 236**a** | 0.028 ± 0.020**b** | -- | 3,009 ± 231**a** |
| Slade et al. 2016ab | DB | 2,342 ± 70 **a** * | -- **a** | 467 ± 38 **a** * | 217,421 ± 11,845 **a** * | -- |
|  | DO | 2,746 ± 220 **a** * | -- **a** | 476 ± 98 **a** * | 235,200 ± 34,306 **a** * | -- |

Differing letters represent significant differences (p<0.05) between groups.
Piccini et al. 2017 not shown since only 1 of 6 dung beetle groups decreased CO<sub>2</sub>e.
\* Data was manually entered, where CH<sub>4</sub> (=25 CO<sub>2</sub>e), CO<sub>2</sub> (=1 CO<sub>2</sub>e), and N<sub>2</sub>O (=298 CO<sub>2</sub>e). CO<sub>2</sub>e represents the sum of CH<sub>4</sub>, CO<sub>2</sub>, and N<sub>2</sub>O;
CH<sub>4</sub>+N<sub>2</sub>Oe represents the sum of CH<sub>4</sub> and N<sub>2</sub>O.

Less than half of current studies reported total CO_2_e (Penttilä et al. 2013, Piccini et al. 2017, Fowler et al. 2020c) which unintentionally obscures an important component: dung beetles’ overall atmospheric effect (Yokoyama et al. 1991ab, Iwasa et al. 2015, Hammer et al. 2016a, Slade et al. 2016b, Evans et al. 2019). Reporting CO_2_e can inform funding agencies which ecological projects may bring about the most benefit – so if CO_2_e is unchanged, then climate change remains unmitigated and potentially no additional C- and/or N-based resources are stored. To fully calculate CO_2_e we suggest not only focusing on CH_4_ and N_2_O emissions (e.g. Slade et al. 2016a) but also aerobic (CO_2_-based) phenomena as this can dramatically alter results (i.e. Iwasa et al. 2015, Evans et al. 2019). So, while studying individual GHGs will still help inform resource recycling mechanisms, such as those found during microbial (Yokoyama et al. 1991a, Slade et al. 2016b) and chemical surveys (Yokoyama et al. 1991ab, Evans et al. 2019); including a net atmospheric effect provides more information at no cost. Therefore, we have complied and analyzed the literature to reevaluate dung beetles’ aggregate GHG effects, report conflicting patterns, and note interesting analyses.

Early research showed that dung beetle groups reduced CH_4_, increased N_2_O, but saw no effect on CO_2_ or CO_2_e when reviewing their pairwise comparisons (see Penttilä et al. 2013 from Table 1) instead of their general ANOVA’s. They theorized that dung beetle activity could reduce anaerobic-based GHGs, primarily methanogenesis (CH_4_ production), by supporting aerobic activity associated with low dung moisture and increased aeration (Penttilä et al. 2013). Meanwhile, Yokoyama et al. (1991ab) showed dung beetles also increased N-release by promoting ammonification, nitrification, and (incomplete) denitrification, thus generating the anaerobic-based N_2_O (Sylvia et al. 2005). Yet if dung beetles reduce CH_4_, but increase N_2_O, and no CO_2_e effect is observed, then are dung beetles climate neutral? By delving into the published literature, we see that dung beetles show no CO_2_e reductions and argue their known effect is negligible (Table 1).

Slade et al. (2016a) reported that dung beetles reduced CH_4_+N_2_Oe (Fig 3 from Slade et al. 2016a) by reducing CH_4_, but not N_2_O, via citing significant treatment by day interactions; however, interactions indicate that treatments were changing differently over time rather than showing overall main effect differences between treatments, which is a subtle, but important, distinction. Our goal is to see if dung beetles can decay dung and reduce dung-based GHGs faster than time alone, which requires a stand-alone treatment effect. Using the reported standard errors, the published data (Supp. Mat. from Slade et al. 2016a) show dung beetles appeared to significantly reduce CH_4_ (dung beetles: 39.8 ± 1.22 vs. dung-only: 46.5 ± 3.73 mg gas/m^2^/d), but not N_2_O (dung beetles: 7.91 ± 0.641 vs. dung-only: 8.07 ± 1.66 mg gas/m^2^/d), leading to no overall CH_4_+N_2_Oe effect (dung beetles: 3,709 ± 210 vs. dung-only: 3,986 ± 581 mg gas/m^2^/d). Understandably, non-overlapping standard errors usually suggest significant differences, but this rule is assumed and does not apply (Skidmore and Thompson 2013) when using unequal sample sizes (n=30: dung beetles vs. n=3: dung-only). Therefore, we retrieved the raw data (Slade, personal communication) and saw there was no overall dung beetle effect (see Supp. Table D7 for the statistical details) for CH_4_ (t=-1.75; df=1,32; p=0.23), N_2_O (t=-0.09; df=1,32; p=0.94), or CH_4_+N_2_Oe (t=-0.49; df=1,32; p=0.64), nor did Slade et al. (2016b) show any differences for CO_2_ for which no raw data was available to analyze (Table 1). Therefore, dung beetles are climate and carbon neutral – not positive. By including main effect analyses and by visibly showing and reporting the extrapolated variation graphically (adding upper and lower variation limits to Fig 3 from Slade et al. 2016a) and numerically (mean ± variation) in the main journal body – we can enhance data analysis and suggest interesting, alternative interpretations.

Hammer et al. (2016a) studied how dung beetles influenced and were influenced by dung from cattle, with and without antibiotics. They also measured dung beetles’ impact on GHG emissions. They saw dung beetles decreased CH_4_, increased N_2_O, and had no effect on CO_2_ emissions relative to the non-antibiotic dung-only (Table 1). Interested in the overall warming effects, we also analyzed the supplemental data supplied online (Hammer et al. 2016b) and saw no dung beetle effect (t=1.17; df=1,19; p=0.25: see Supp. Table D7 for statistical details) when comparing non-antibiotic dung pats with and without dung beetles, respectively (Table 1).

Piccini et al. (2017) investigated whether dung beetle diversity affected GHGs differently as our experimental design currently does. The authors favored using the unadjusted p-value rather than the adjusted p-value (which accounts for familywise error and reduces the number of false positives – Wilcox et al. 2013) though both were supplied. Using their reported p-adjusted values for their general ANOVA’s, an alternative interpretation shows that the dung beetles (treatment only) had no cumulative effect on CH_4_, CO_2_, or N_2_O fluxes, though there were CO_2_e differences (Supp. Table F from Piccini et al. 2017). Even so, the modified GLS model (Supp. Table I from Piccini et al. 2017) and multiple pairwise comparison tests (Supp. Table H from Piccini et al. 2017) showed that only the one of the six dung beetle treatment, the most diverse group, reduced CO_2_e. This alternatively suggests that most experimental dung beetle groups are climate neutral.

Some studies found beneficial effects, others found negative effects or conflicting patterns. Iwasa et al. (2015) reported that dung beetles reduced CH_4_, but greatly increased CO_2_, N_2_O, and, when approximated, CO_2_e (Table 1). Evans et al. (2019) found dung beetles increased CO_2_ on specific days and, surprisingly, CH_4_ overall – even though these gases are frequently produced under radically different conditions. Regardless, dung beetles had an increased (climate warming) effect on N_2_O, CH_4_+N_2_Oe (Evans et al. 2019), but not on CO_2_ or CO_2_e when approximated (Table 1). The pattern becomes that Penttilä et al. (2013), Hammer et al. (2016a), Slade et al. (2016ab), Piccini et al. (2017), and Evan et al. (2019) found dung beetle activity to be climate neutral, while Iwasa et al. (2015) found dung beetle activity to be climate negative (i.e. a GHG source).

We initially studied (Supp. Table D1, Supp. Fig G’s) whether different dung beetle activities (tunneling vs. dwelling) under field chamber conditions influenced GHGs differently, but discovered negligible CO_2_e effects. The reexamined literature also suggests this. One methodological aspect common within all of these studies, including our original experiment, is that we homogenized and standardized the dung prior to adding dung beetles – but if mixing (homogenizing) aerates the dung and reduces GHG-producing microbes, what is the difference between mixing the dung by hand or with dung beetles? We hypothesize that: 1) mixing dung obscures a dung beetle’s effect on GHGs, and 2) that dung beetles can reduce CO_2_e. By including a homogenization (unmixed dung-only) control, repeating the experiment, and increasing our replicates, we asked whether the negligible dung beetle effect was because of randomness, methodology, or incomplete theory.

## Materials and methods

### Experimental Design

Here we report *two designs* and perspectives from a *single dataset*: the combined 2016-2017 design (Supp. Table D1, Figs G1-G4) and the 2017 unmixed design (Supp. Table D2, Figs 3-6). The **combined design** asks if dung beetle activity (tunneling vs. dwelling) affects the mixed dung-only using more replicates, while the **unmixed design** asks if mixing (homogenization) itself confounds the results. Since it is unknown whether mixed versus unmixed dung produces different GHG fluxes, dung beetles were added only to the standardized, mixed dung to reduce labor/costs, redundancy, and unwanted assumptions. The results from the combined and unmixed designs are practically identical (see Supp. Section A for minute differences) and so we will focus only on the more extensive unmixed design, which also forms a part of the combined design (see Table 2’s footnotes).

**Table 2.** The experimental designs used in 2016 and 2017

| Treatment | Beetle | Dung | Grass | Purpose |
| --- | --- | --- | --- | --- |
| Tunn | ✓ | ✓ | ✓ | To visualize if different dung behaviors (tunneling vs. dwelling) affects dung generated GHGs in different ways. |
| Dwell | ✓ | ✓ | ✓ |  |
| TunnDwell | ✓ | ✓ | ✓ |  |
| Mixed Dung |  | ✓ | ✓ | The standardized treatment controls allow for comparison and environmental monitoring. |
| Grass |  |  | ✓ |  |
| Unmixed Dung |  | ✓ | ✓ | The homogenization controls assess if this methodology confounds results. |
| Pasture |  |  | ✓ |  |

Where Tunn (=Tunneler, *Onthophagus taurus*) and Dwell (=Dweller, *Labarrus pseudolivinus*)
Combined design (=Standard treatments only from both 2016 & 2017); n=21 rep/treatment/day.
Unmixed design (=Pasture + Standard treatments from 2017 only); n=12 rep/treatment/day.
Vegetative Controls = Pasture or Grass-only treatments.
Unmixed Dung = unhomogenized, unstandardized, untouched, variable.
Mixed Dung = homogenized (mixed), standardized, set amount.

We repeated our experiments once per month (n=4 experiments) during the summer totaling 12 replicates per treatment for the unmixed design (June to September 2017) and 21 replicates for the combined design (May to September 2016, June to September 2017). Using a randomized complete block design (n=3 blocks/experiment), we measured GHGs (CH_4_, CO_2_, N_2_O) over the course of two weeks (0, 1, 3, 7, 14d) using GHG chambers (n=1 treatment/block; Fig 1) at the grassy pastures of the beef unit (Lat. 35°43’47.40”N, Long. 78°41’15.50”W) at NCSU Lake Wheeler Road Field Lab (Raleigh, North Carolina, USA). On both 0d and 14d, we: 1) collected and dried dung and soil cores in the oven at 55°C to measure moisture loss (Supp. Table D6; for both data and method descriptions), and 2) visually monitored dung damage (see photographs in Fig 2) to track dung beetle-induced abiotic changes. The dataset included a variety of pasture sites, weather conditions, dung compositions, and dung beetle populations as is naturally expected when replicating across various seasons, years, and herds, and this helps increase scientific rigor and applicability (Casler 2015).

**Fig 1.**
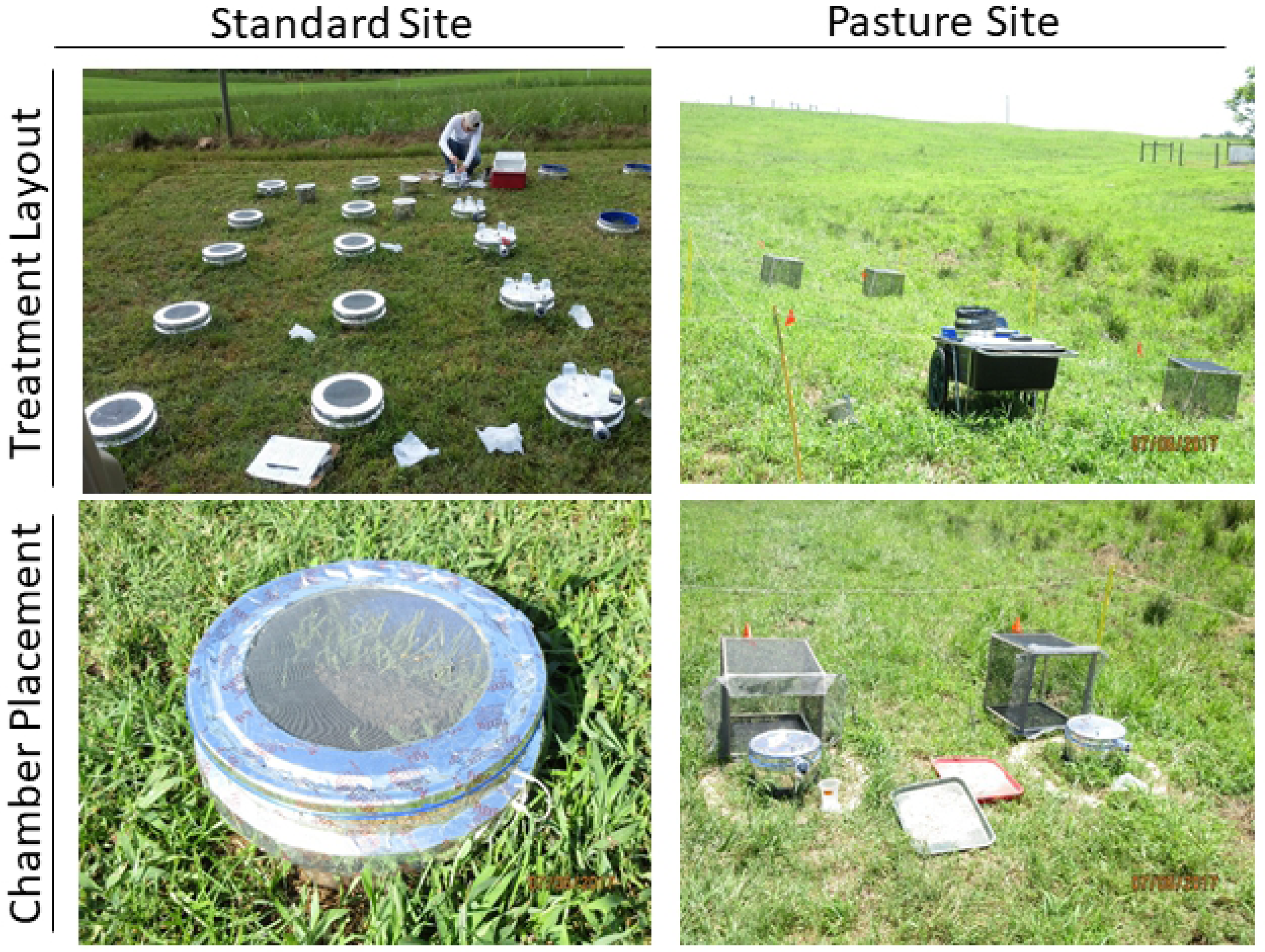
Layout and chamber positioning. The standard site consists of a mowed field outside of cattle pastures, while the pasture site consists of a recently used cattle pasture. See Supp. Section A for more detail about site differences.

**Fig 2.**
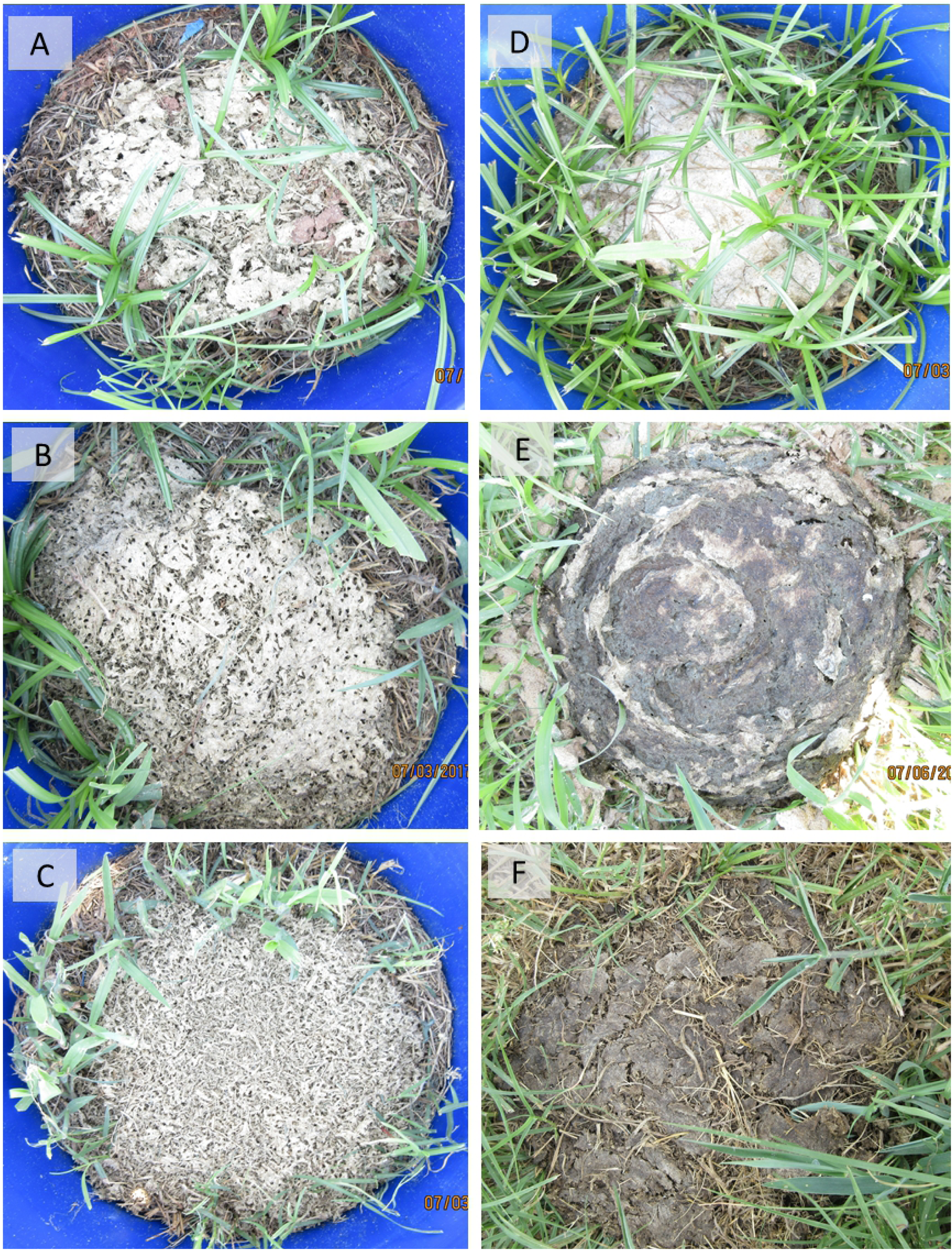
Visual comparison of experimental treatments. Examples of the (A) Tunn; (B) Dwell; (C) TunnDwell; (D) Mixed Dung; and (E) Unmixed Dung treatments; including a naturally colonized dung pat (F). Similarities between the dung beetle occupied treatments and the native dung pat suggests study treatments were representative of natural dung beetle activity.

**Fig 3.**
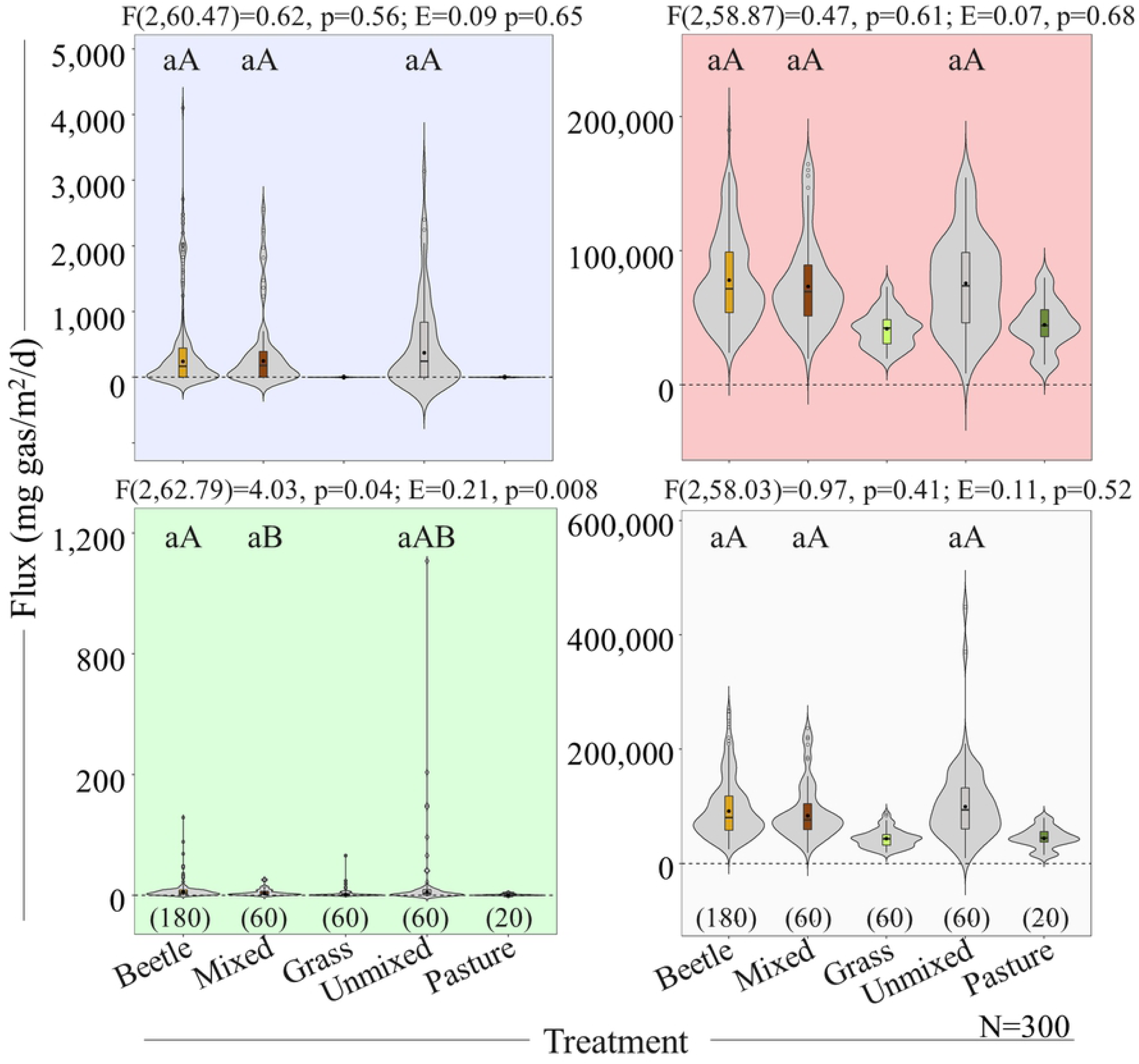
Violin box plots of HMR Flux by Treatment. Each quadrant represents a GHG including CH_4_ (top left), CO_2_ (top right), N_2_O (bottom left), and CO_2_e (bottom right) with their respective omnibus mean-based ANOVA’s (F_df num, df den_) and median-based Explanatory Effect (E) Sizes shown above. Pairwise comparisons of ANOVA’s (lowercase letters) are shown within the graph. Sample sizes (n) are shown underneath each treatment and total samples (N) are shown along the x-axis. Differing letters between groups show differences (p≤0.05). Exact means, medians, and measures of variations are found in Supp. Table D2.

**Fig 4.**
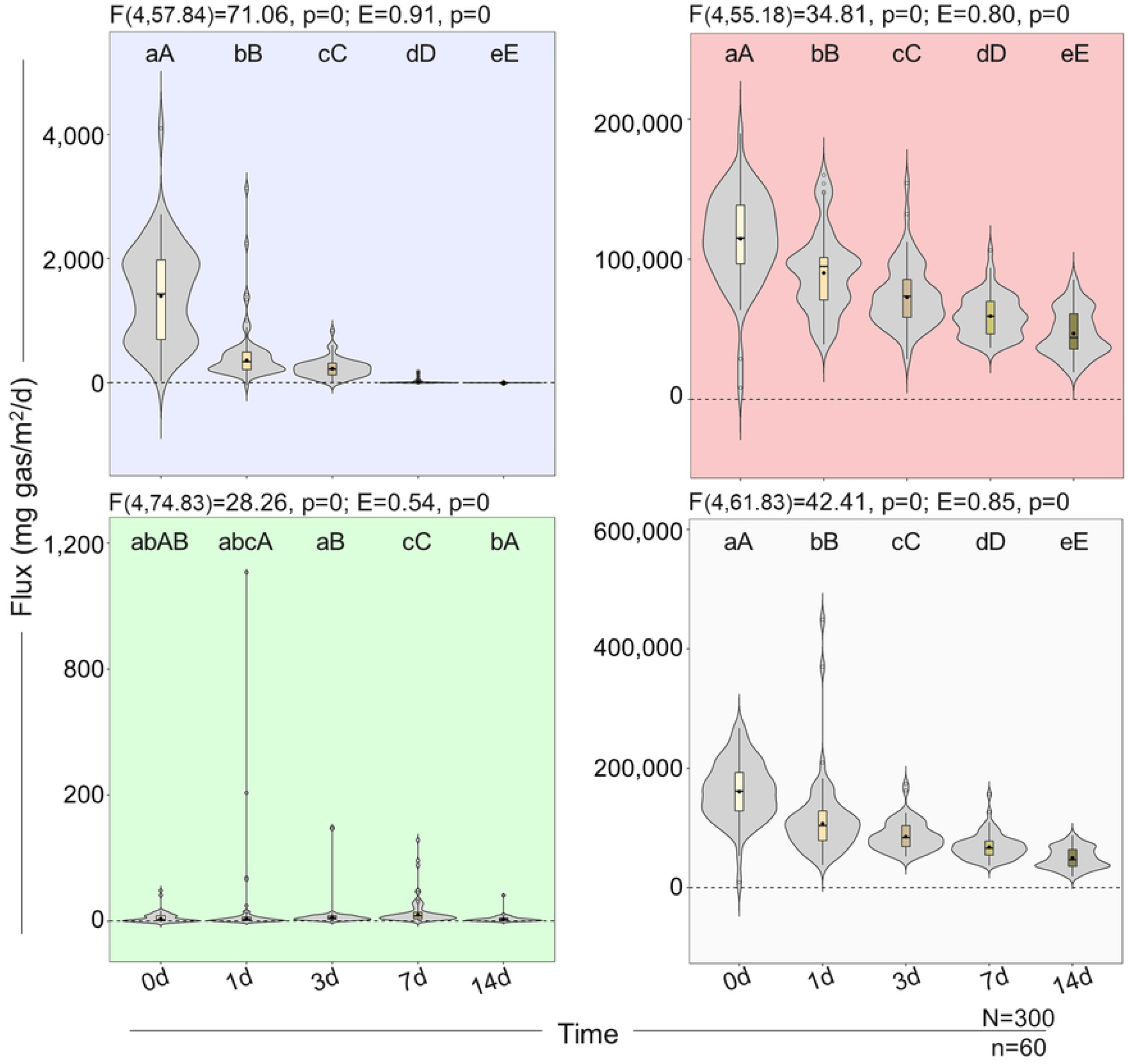
Violin box plots of HMR Flux by Time. Each quadrant represents a GHG including CH_4_ (top left), CO_2_ (top right), N_2_O (bottom left), and CO_2_e (bottom right) with their respective omnibus mean-based ANOVA’s (F_df num, df den_) and median-based Explanatory Effect (E) Sizes shown above. Pairwise comparisons of ANOVA’s (lowercase letters) and Effect Sizes (uppercase letters) are shown within the graph. Sample size (n) and total samples (N) are shown along the x-axis. Differing letters between groups show differences (p≤0.05). Exact means, medians, and measures of variations are found in Supp. Table D2.

**Fig 5.**
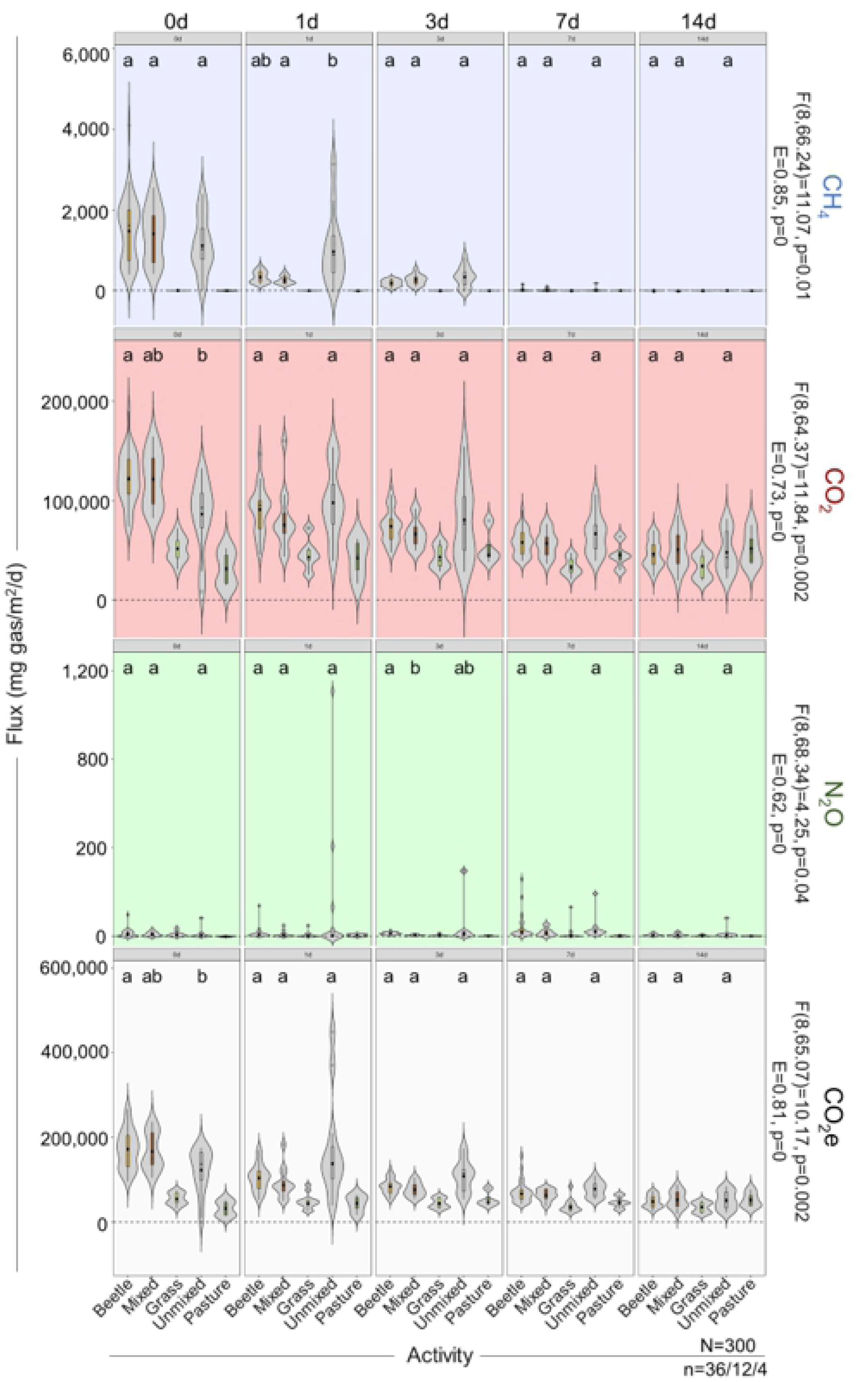
Violin box plots of HMR Flux by Treatment within Time. Each quadrant represents a GHG including CH_4_ (top left), CO_2_ (top right), N_2_O (bottom left), and CO_2_e (bottom right) with their respective omnibus mean-based ANOVA’s (F_df num, df den_) and median-based Explanatory Effect (E) Sizes shown above. Pairwise comparisons of ANOVA’s (lowercase letters) are shown within the graph. Sample sizes (n) are shown underneath each treatment and total samples (N) are shown along the x-axis. Differing letters between groups show differences (p≤0.05). Exact means, medians, and measures of variations are found in Supp. Table D2.

**Fig 6.**
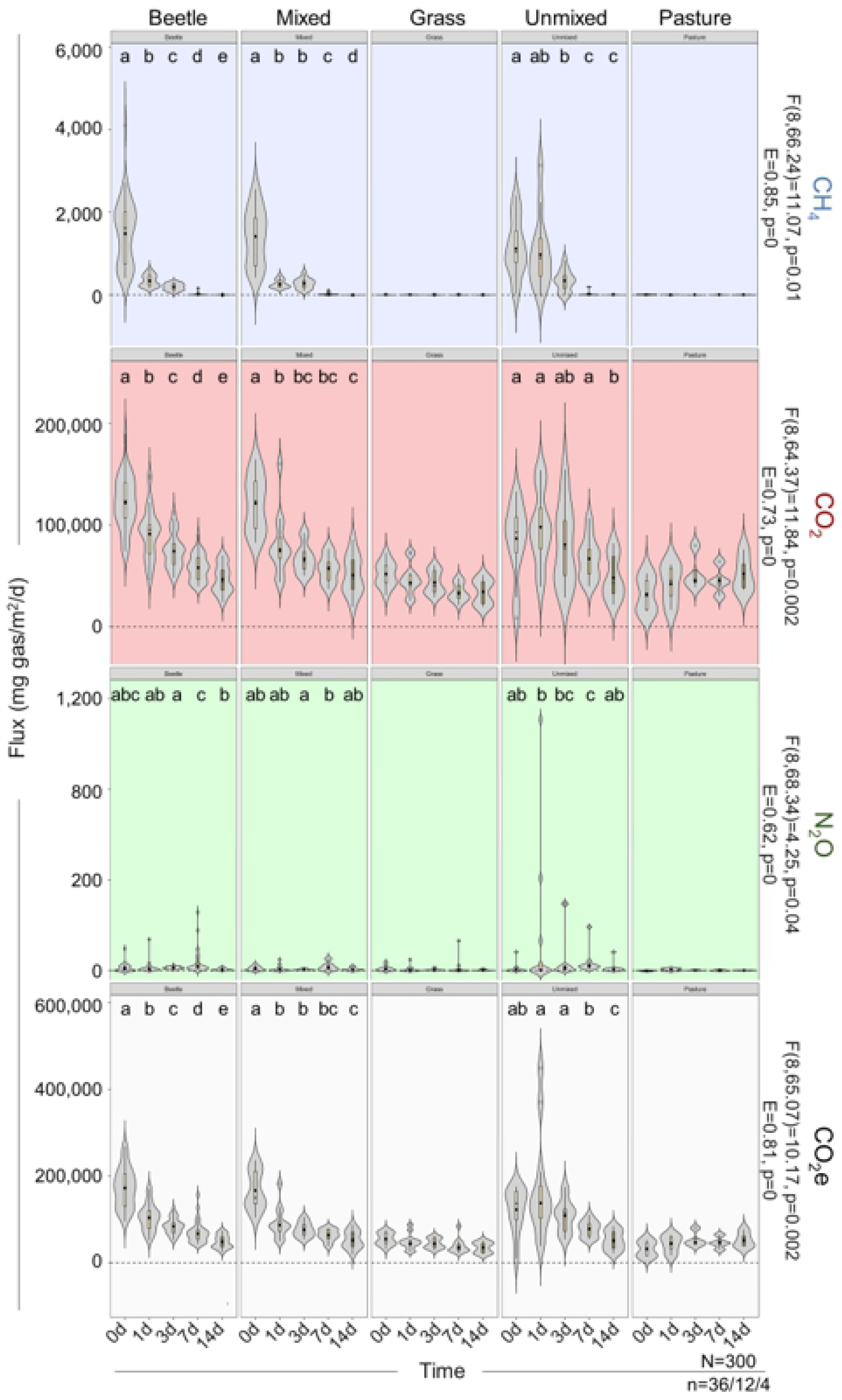
Violin box plots of HMR Flux by Time within Treatment. Each quadrant represents a GHG including CH_4_ (top left), CO_2_ (top right), N_2_O (bottom left), and CO_2_e (bottom right) with their respective omnibus mean-based ANOVA’s (F_df num, df den_) and median-based Explanatory Effect (E) Sizes shown above. Pairwise comparisons of ANOVA’s (lowercase letters) are shown within the graph. Sample sizes (n) are shown underneath each treatment and total samples (N) are shown along the x-axis. Differing letters between groups show differences (p≤0.05). Exact means, medians, and measures of variations are found in Supp. Table D2.

The unmixed design called for two distinct sampling locations and treatments (Table 2) using cattle dung from the same beef herd: the **standard** site and the **pasture** site (Fig 1) of which detailed differences are described in Supp. Section A. The standard site required we bury GHG chambers outside of cattle pastures, such that the anchor wall was five inches tall, at least 3 weeks pre-experiment to mimic published designs (e.g. Penttilä et al. 2013, Slade et al. 2016a). On 0d, we added ∼1000g of fresh (<5min old), premixed (by hand, dung only) cattle dung (Supp. Fig A) and the appropriate treatments. Meanwhile, the pasture site measured arbitrarily lain dung pats within cattle pastures using mobile GHG chambers alongside netted cages (Fig 1), which: a) avoided chamber-induced microclimates, b) reduced destructive chamber burial on pastures, andc) prevented dung arthropod entry. Both sites used the same sampling equipment (Fig 2). Fowler et al. (2020b) discusses and analyzes the physical chamber designs and the gas sampling strategies.

### Dung Beetle Collection & Treatment Layout

Dung beetles were grouped by nesting behaviors: endocoprids (“dwellers” or Dwell), paracoprids (“tunnelers” or Tunn), and telecoprids (“rollers”). Dwellers ‘shred’ or ‘mix’ the dung from within; tunnelers bury dung (‘brood’) balls beneath the pat; rollers roll the dung ball away and bury it elsewhere (Bertone et al. 2004). We focused specifically on dwellers and tunnelers because: 1) In North Carolina, the tunneler, *Onthophagus taurus* (Schreber), and dweller, *Labarrus* (*Aphodius*) *pseudolividus* (Balthasar), are abundant on cattle pastures (Bertone et al. 2005) – this ensured adequate replication across seasons and years; 2) both tunnelers and rollers exhibit burial activity, thus studying dwellers and tunnelers fully represent our desired behavioral repertoire. Following an additive design, we measured each dung-use group together and apart, while also including mixed dung-only, unmixed dung-only, and grass/pasture-only (‘vegetative’) controls (Table 2). Labor and material constraints restricted additional replicates solely for the pasture-only control (n=4), but this unbalanced design was accounted for statistically by using more conservative analyses (see *Statistics*). We collected dung beetles by floating and sorting them (see Fowler et al. 2020a for methodology, stats, and other details) within 48 hrs pre-experiment and held them in incubators at 12°C (L:D 16:8) with moistened towelettes until use. We measured 3 grams of dung beetles^3^ per treatment (Table 2) to avoid confounding dung beetle size and number. This biomass was selected because the damaged dung of our treatments (Fig 2A-C) was similar to the dung damage found naturally in the field (Fig 2F), which indicated an optimal representation of dung beetle activity. Additionally, we conducted small-scale respiration studies pre-0d to examine if dung beetles produced ample greenhouse gases, but dung beetles were found only to respire elevated CO_2_ and so will only be briefly discussed (see Supp. Section B for more detailed methods and analyses).

### Statistics

First, we conducted power tests (packages: “pwr”, “pwr2”) in the R statistical program (R Development Team, Geneva, Switzerland; http://www.r-project.org) to estimate the required sample sizes for 60% power given our number of contrasts (Supp. Mat. P). Second, we acknowledged any heterogeneous variation, extreme outliers, and positive skewness (Erceg-Hurn and Mirosevich 2008) using Wilcox’s Robust Statistics (package: WRS) (Wilcox 2013) by:

A. Winsorizing extreme outliers to allow focus on gas majority representation,
B. identifying skewed outliers using modified M-estimators,
C. bootstrapping the data (n_boot_=500 to 600) to calculate the proportion showing p≤0.05, and
D. using the Benjamini–Hochberg procedure to account for familywise error, which creates Type II errors (“false positives”) by chance when evaluating multiple comparisons.

We did this for both mean-based (ANOVA) and median-based (Effect Size) analyses for both the combined (Supp. Table D1, Supp. Figs G1-G4) and unmixed designs (Supp. Table D2, Figs 3-6). Lastly, we provided a simplified treatment layout (Figs 3-6) which combined all dung beetle treatments (Tunn, Dwell, and Tunn + Dwell otherwise known as TunnDwell) into a single group (Beetle) for ease-of-use since the results were identical (Supp. Table D1 vs. Supp. Table D2).

### Conversions, Tables, & Graphs

We converted our original GHG measurements (ppm) into fluxes (mg gas/m^2^/d) using sampling time, headspace volume (Chamber Volume - Dung Volume), and temperature expansion ratios, and by applying the modified Hutchinson and Mosier method (package: HMR) to calculate linear/nonlinear HMR fluxes (Venterea and Parkin 2012). Total CO_2_e were calculated as the sum of the global warming potential impact factor over 100 years (IPCC 2007) as follows: CH_4_ (=25 CO_2_e), CO_2_ (=1 CO_2_e), and N_2_O (=298 CO_2_e). We have included treatment, time, and treatment:time interactions (marginal means), graphs, and tables for both the combined design (Supp. Figs G1-4, Supp. Table D1) and unmixed design (Figs 3-6, Supp. Table D2). Relevant statistics (ANOVA’s + effect sizes) are presented on all figures, but we also provided supplemental statistics (Supp. Section D), power analyses (Supp. Section P), respiratory analyses (Supp. Section B), and graphs (Supp. Section G).

## Results

By using the unmixed design (Figs 3-6) we tested if mixing obscured a dung beetle effect, and by using the combined design (Supp. Figs G1-4) we tested if dung beetles had any effect. Any treatment differences reported here are not due to site or vegetative (Grass-only vs. Pasture-only) differences as we found no overall vegetative differences for CH_4_ (t=0.05, p=0.96), CO_2_ (t=0.99, p=0.33), N_2_O (t=1.98, p=0.054), or CO_2_e (t=0.82, p=0.42) nor dung-to-chamber volume ratios between sites (Supp. Fig G6). We will examine two types of statistics here to present a more nuanced view of our data: the p-value and effect sizes. The p-value is the likelihood that the between-group mean difference (≠0) seen is potentially a false positive a certain percentage (∼5% when α=0.05) of the time given accurate ANOVA assumptions and sufficient power (Colquhoun 2014). Meanwhile the effect *size* reflects the *magnitude* of reported differences to help aid interpretative conclusions (a small/weak, medium, and large/strong explanatory measure of effect is considered at least E=0.15, 0.30, and 0.50, respectively – Wilcox et al. 2012).

### Methane

Treatment had no effect on CH_4_ fluxes in the combined (E=0.008, Supp. Fig G1) or unmixed design (E=0.09, Fig 3); meanwhile time steadily reduced CH_4_ by >200x by 7d (E=0.91-0.98, Figs 4, G2) showing time as the strongest predictor. Only TunnDwell groups reduced CH_4_ by 2.31x compared with mixed dung-only on 3d alone (Supp. Fig G3). Similarly, unmixed dung-only produced 2.83x and 3.71x more CH_4_ (Supp. Table D2) than dung beetles and mixed dung-only groups (Fig 5) solely on 1d – revealing that neither dung beetles nor hand-mixing meaningfully affected weekly CH_4_ fluxes compared to natural decay. Although the unmixed dung-only produced 1.50x (mean) and 1.38x (median) more than the mixed dung-containing treatments overall (Supp. Table D2), the general effect was negligible (Fig 3). This increase could be because:

1. *site-based vegetative differences*: however, the grass-only and pasture-only controls (Table 2) produced <1 mg CH_4_/m_2_/d on any given day, so all CH_4_ fluxes are dung-based;
2. *different chamber and dung volumes between sites*: this could theoretically bias certain treatments toward greater fluxes, but given that unmixed dung-only sizes were random (often larger) and measured using field-based chamber methods (larger volume) - the total dung-to-chamber volume ratios between sites were similar (see Supp. Section A for a more in-depth discussion of calculations and analyses). After all, any biases would show up on 0d if the ratios were different, but this did not occur (Supp. Fig G6); and
3. *between-site variances*: mixed (homogenized) dung in the semi-permanent (buried) chambers could have reduced variation compared with unmixed dung (non-homogenized) of the non-standardized (unburied) pastured treatments, thus resulting in reduced power to detect differences. Yet no variation differences were found between the unmixed and mixed dung-only (p>0.05) across a week (Supp. Fig G8).

If mixing itself was a factor in affecting GHGs we would expect to see differences between the mixed and unmixed dung-only treatments on 0d. However, there were no differences and the mixed dung-only treatment produced slightly more CH_4_ despite being presumably more aerated. Likely it is because fresh cattle dung is liquid-like, and so mixed dung easily reforms and inhibits aeration. However, dung beetles physically affected the dung (Fig 2) through more constant mixing, enhanced desiccation (Supp. Table D6), and presumably greater aeration; yet, we observed no dung beetle advantage in reducing CH_4_, thereby suggesting time and dung-presence are the only meaningful factors for CH_4_ generation.

### Carbon Dioxide

Treatment had no effect on CO_2_ fluxes in either the combined (Supp. Fig G1) or unmixed design (Fig 3). Comparatively, time had a strong reductive effect (E=0.70-0.80) compared with treatments (E=0.07-0.10), as CO_2_ fluxes declined by >1.9x from 0d to 7d (Supp. Fig G2) or 14d (Fig 4). Oddly, we expected dung beetles to increase CO_2_ fluxes given their own respiration and aerating/aerobic-based activities, but our (Supp. Fig B) and Piccini et al. (2017)’s respiration studies showed that dung beetle respiration was <1.5% of a single day’s CO_2_ emissions (see Supp. Section B for further discussion). Curiously, while treatments mostly showed no differences (Supp. Fig G3), the unmixed dung-only produced less CO_2_ fluxes than the mixed dung groups on 0d (Fig 5) and showed slower week-long declines (Fig 6). This was, respectively, due to the pasture-control influencing the unmixed dung-only’s reported fluxes, for example:

1. by producing 1.62x less CO_2_ (t=0.19, p=0.08) than the grass-only on 0d (Supp. Table D2), and
2. by producing 1.07x more CO_2_ (t=2.41, p=0.056) over time (Fig 6) because it was not regularly cut as the grass-only was and grew over time. Thus, the drop in CO_2_ for the unmixed dung-only group was similar to other mixed dung groups and was masked initially, thus the decline appeared more slowly – regardless, any small differences were negligible (p>0.05).

In all, dung-containing groups produced >1.69x the CO_2_ compared with their vegetative controls, thus showing dung-presence enhances CO_2_. Overall, dung beetles had no CO_2_ effect, while only time and dung-presence had strong, reliable effects (Fig 4).

### Nitrous Oxide

Unlike the other gases, treatment showed a small-to-medium effect on median N_2_O fluxes for both the unmixed design (E=0.21, Fig 3) and when combining years (E=0.30, Supp. Fig G1), while time had a small-to-strong effect (E=0.22, Supp. Fig G2; E=0.54, Fig 4). As with all main effect analyses, data aggregated across time or treatment obscures differences between high-performance treatments and strong time effects, therefore day-by-day analysis was required for differentiation. For the treatments: the unmixed design showed dung beetle groups producing an average of 2.6x more N_2_O than the mixed dung-only on 3d alone (Fig 5, E=0.62). This increase was due to the Tunn and TunnDwell groups producing >2.6x and >1.86x the average N_2_O of the mixed dung-only and Dwell, respectively (Supp. Fig G3, E=0.44). This suggests specific dung beetle behavior matters. Interestingly, while there were no variation differences (p>0.05) between the unmixed and mixed dung-only (Supp. Fig G8), the dung beetle groups produced a greater frequency of larger fluxes (p=0) and a smaller frequency of minimums (p=0) (Supp. Fig G7) reflecting the significant omnibus and effect size analyses (Fig 3). This suggests tunneler-activity, but not mixing nor dwelling-activity, specifically generated more N_2_O (1.57x) than unmixed dung-only despite its weak effects (Fig 3). Curiously, our vegetative controls did not show N_2_O spikes following rain events, as they do in agricultural soils (Sylvia et al. 2005), perhaps because vegetation retains moisture more consistently across time than bare soil, and so produces consistent low-emissions in grassy pastures.

Nevertheless, the strongest effects on N_2_O were time and dung-presence, especially relative to the vegetation-only controls. Grass and pasture-only treatments produced similar, low-emission trends across time (Supp. Table D2) (Fig 6), often generating 0.10-0.50x the amount of the dung-containing treatments on any given day. Across time, the dung-containing groups produced 2.59-7.26x more N_2_O than their respective vegetative controls, thus showing dung’s propensity for N_2_O generation (Fig 6) until complete decay (see the desiccated dung in Fig 2). Dung (Supp. Fig G9) and soil (Supp. Fig G10) moisture loss regressions showed that greater soil moisture was positively correlated (R^2^=0.33, p=0) with greater N_2_O fluxes as expected, but there was surprisingly no relationship with dung moisture content (R^2^=0.02, p=0.98) – in contrast, CH_4_, CO_2_, and CO_2_e were positively correlated with dung moisture loss (Supp. Fig G9, p≤0.01), but not soil moisture loss (Supp. Fig G10, p≥0.05), despite the positive correlations between soil and dung moisture loss. Understandable given that CH_4_ is generated almost solely by dung (dung-presence generated 622.8x more CH_4_ than vegetation-only), while CO_2_/CO_2_e is heavily influenced by dung moisture simply because it is also correlated with dung decay, gradual aeration, and other time-related variables (multicollinearity effect). But N_2_O offers a different picture: dung-containing treatments held 1.22x more soil moisture than the vegetation-only (Supp. Table D6), with Tunnelers reducing dung moisture content by 1.78x compared with mixed dung-only (Supp. Table D6, p=0). In short, drier dung correlated with increased soil moisture, likely from dung leaching both moisture and nutrients to the surrounding soil, with active dung-burial species potentially enhancing this process and increasing their N_2_O fluxes due to soil activity (Supp. Fig G3).

Each quadrant represents a GHG including CH_4_ (top left), CO_2_ (top right), N_2_O (bottom left), and CO_2_e (bottom right) with their respective omnibus mean-based ANOVA’s (F_df num, df den_) and median-based Explanatory Effect (E) Sizes shown above. Pairwise comparisons of ANOVA’s (lowercase letters) and Effect Sizes (uppercase letters) are shown within the graph. Sample size (n) and total samples (N) are shown along the x-axis. Differing letters between groups show differences (p≤0.05). Exact means, medians, and measures of variations are found in Supp. Table D2.

### Total Greenhouse Gas Effect

CO_2_e is calculated by determining the relative effects of each GHG. For example, 98.87% of all GHGs collected was solely CO_2_, but since CH_4_ and N_2_O enjoy a larger greenhouse effect, they respectively contributed to 23.07 and 7.28% of the total effect (Supp. Fig G5). Even so, CO_2_ commands 69.65% of the sum-total CO_2_e which is why CO_2_e graphs (Figs 3-6, Supp. Figs G1-G4) predominately follow CO_2_ trends. Treatment had no effect (∼1x) on CO_2_e (E<0.12, Fig 3), and the small reduction of CO_2_e on 0d was attributed to the vegetative differences as described in the *Carbon dioxide* section. Comparatively, time steadily reduced CO_2_e by 3.24x (Fig 6) over the course of a week (E=0.85, Fig 4). Ultimately, dung beetles were climate neutral and carbon neutral (neither storing nor releasing resources).

In summary, we see that neither dung beetles nor mixing affected the average CH_4_, CO_2_, N_2_O, and CO_2_e, though dung beetles did increase N_2_O medians. Time, meanwhile, drastically reduced CH_4_, CO_2_ and CO_2_e across a week, while inconsistently increasing N_2_O resulting from dung beetle interactions. Ultimately, we eliminated our hypothesis that mixing masked dung beetle effects and found no dung beetle effect on GHGs.

## Discussion

Since the early 1980s, an increasing number of researchers studied how arthropods and annelids – such as termites (Sugimoto et al. 2000), dung beetles (Yokoyama et al. 1991ab), and earthworms (Lubbers et al. 2013) – affected the GHG emissions of plant litter, soil, and/or dung. These arthropods affected the decomposition rates and microbial pathways driving carbon (C) and nitrogen (N) storage/release throughout the environment. For example, by reducing C and N lost to the atmosphere, it is instead used and stored terrestrially (Sylvia et al. 2005). Theoretically, dung beetles are capable of similarly affecting GHGs, but lacked supportive research (see *Introduction*). By improving the power of our study (combined-years design) and testing potential methodological problems (unmixed design), we suggest that dung beetles are ultimately carbon neutral and that the physical ‘mixing’ of dung may not be a significant mechanism in reducing GHGs.

### GHG trends and their potential causes

As the pat ages, the constant decay physically alters the dung by leaching/evaporating water (by 1.89x from 0-14d, Supp. Table D6) and loosening the dung structure – a process aided by the disturbance and disassembly of dung pats by dung beetles (Fig 2). The disintegrating and desiccating dung allows for deeper oxygen (O_2_) penetration and permeation such as seen in soil (Sylvia et al. 2005). If true, we would predict decreased CH_4_, increased N_2_O, and increased CO_2_ over time. However, we saw both dung-based CH_4_ and CO_2_ (and so CO_2_e) decline permanently over time until they mirrored the vegetative-control fluxes (Fig 6). Though expected for CH_4_, CO_2_’s decline was a surprise. Presumably transitioning from an anaerobic to an aerobic dung pat by mixing or aging should predictably increase CO_2_ emissions via environmental respiration or enhanced gas transport, though not dung beetle respiration (Iwasa et al. 2015). After all, we (Supp. Section B) and Piccini et al. (2017) showed that dung beetle respiration was less <1.5% of the total CO_2_, and that dung beetles did not release more CH_4_ and N_2_O than the control (Supp. Fig B), despite consuming methanogen and denitrifier-rich dung (Yokoyama et al. 1991a).

Ultimately, no CO_2_ differences existed between any treatments (Fig 3) even after two weeks of decomposition and desiccation (Fig 2). Meanwhile, N_2_O followed our predictions but offered a surprise.

Generally, N_2_O emissions result from incomplete byproducts of denitrification (Sylvia et al. 2005), nitrate reduction (Penttilä et al. 2013, Slade et al. 2016a, Piccini et al. 2017), nitrification (Iwasa et al. 2015), from increased microbial abundance and activity (Yokoyama et al. 1991ab), and/or increased gas transport – such as when dung beetles microtunnel into wet dung (Evans et al. 2019). However, the greatest (>90%) N_2_O production is formed by incomplete denitrification, when:

1. there is sufficient O_2_-disrupting amounts – too little O_2_ forms the benign, atmospheric N_2_; too much O_2_ forms the pre-GHG cursor, NOx; and just the right amount forms N_2_O due to oxygen-inhibited microbial enzymes (Sylvia et al. 2005). O_2_ competes with water to fill soil pores, and so a substrate’s moisture can inhibit aeration – hence why N_2_O emissions spike after rainfall (Sylvia et al. 2005);
2. there are large NO3 ^-^ pools (akin to synthetic nitrate fertilizers – Akiyama et al. 2010) that microbes use to produce N_2_O. Dung beetle activity provide NO3 ^-^ pools by enhancing ammonification and nitrification through aerobic soil activity (Yokoyama et al. 1991b). Collectively this suggests that cow dung is an obvious moisture and fertilizer source, and that dung beetles may increase the incomplete denitrification rate.

The increased N_2_O fluxes from 1-7d and the sharp 14d decline in dung-based treatments suggest that our treatments supported large enough N-pools and a mostly anaerobic state sufficient for N_2_O production until 14d. However, we wondered if dung or soil was the main source of denitrifier activity. Consider that: higher soil moisture, but not dung moisture (Supp. Fig G9), predicted higher N_2_O emissions (Supp. Fig G10) despite dung and soil moisture content being correlated (r=0.36 to 0.64 depending on the gas, Supp. Table D8). This suggests that dung leaches moisture and resource-rich fluids to the surrounding soil and so soil microbes may be generating N_2_O. In support, research shows that both C and N-based water-soluble nutrients increased with dung-presence (Yokoyama et al. 1991a, Evans et al. 2019) and all soil-based, N-acting bacterial and fungal groups (Yokoyama et al. 1991a) increased in dung-containing groups. Thus, soil microbes possess all the necessary prerequisites for N_2_O generation. We also see that dung-only treatments possessed higher soil and dung moisture contents (Supp. Table D6) resulting in treatments growing moisture-loving white fungus or mushrooms, but curiously also had lower N_2_O fluxes than the tunneler-containing groups that sported lower dung soil moisture contents (Supp. Table D6). Combined, this suggests that though tunneler burial activity decreases dung and soil moisture through churning – the soil disturbance, dung incorporation, and aeration likely generate more N_2_O than dung-presence alone.

In short, we suggest the higher N_2_O generation associated with dung-presence is likely due to soil-generation. The dung provides the fertilizer, the moisture or rain leach the dung-rich resources, and any additional soil churning, particularly by the burial-heavy tunneler groups, aerate and trap dung-resources for greater microbial consumption. After all, dung beetles increase C, N, and other soil analytes (Nichols et al. 2008), and although resource-deposition does not necessarily coincide with exact GHG production dates (Evans et al. 2019), these materials likely remain available until microbial-digestion conditions are ideal. Altogether, dung-presence, increased soil moisture, and tunneler-presence each increased soil-based N_2_O fluxes. These processes mirror common N_2_O spiking agriculture practices such as: applied fertilizers (Akiyama et al. 2010), rain-filled soils (Sylvia et al. 2005), farmer pre-rain fertilizer applications (Singh and Sekhon 1979), churning compost (Lim et al. 2016), and tilling soils (Snyder and Hendrix 2008).

### Comparing Dung Beetle Activity

Through a literature review, we sought to uncover if the dung beetle effect was consistent despite a diversity of methods, environmental conditions, sample sizes, treatments, dung beetle activity/biomass, and statistical focuses (Table 4).

#### Methane

Of our combined studies: 3/7’s presumed dung beetles decreased CH_4_ over time because of aeration, 3/7’s showed no effect, and 1/7th presumed dung beetles increased CH_4_ when they created alternative gas pathways (microtunnels) in wet dung. Periodic CH_4_ increases are not uncommon (Penttilä et al. 2013, Slade et al. 2016a, Evans et al. 2019) and highly suggest methanogen-preferred anaerobic conditions attributed to wet conditions. Thus, as pats dried, most studies showed extinguished CH_4_ fluxes by 7d (Piccini et al. 2017, Fowler et al. 2020c) or 30d (Penttilä et al. 2013, Slade et al. 2016a). Even Evan’s et al. (2019)’s study showed decreased dung-generated CH_4_ relative to the pasture-only by 7d despite the meadow being a strong CH_4_ sink.

#### Carbon dioxide

Of our combined studies: 1/7th presumed dung beetles increased CO_2_ flux from either dung-beetle respiration (Iwasa et al. 2015) or enhanced gas transport on particular days (Evans et al. 2019), while 6/7’s showed no effect (Table 4). Interestingly all dung-only controls followed negative distributions (linear, exponential, Gaussian) for CO_2_ over time – suggesting decreased biological respiration is very common unless enhanced by other factors (e.g. dung beetles; Iwasa et al. 2015, Evans et al. 2019).

#### Nitrous oxide

Of our combined studies: 4/7’s of studies showed no dung beetle effect (including ours, though we showed a median increase), while 3/7’s showed large N_2_O increases based on numerous explanations. Like CH_4_, most dung-containing treatments saw periodic increases, except for Piccini et al. (2017). This suggests that dung-presence and dung-beetle presence can spike N_2_O fluxes. Interestingly, in every study where CH_4_ was significantly influenced, so was N_2_O.

Despite this information, time is the strongest predictor in all studies and consistently decreased CH_4_ and CO_2_’s emissions and increased N_2_O’s emissions (Table 3).

**Table 3.** A summary of reported GHG flux differences (+: increase, -: decrease, 0: no effect) and their overall trajectory surrounding aggregate time effects of ANOVAs (p<0.05) of the dung-containing treatments (DC), or on the dung-only treatments (DO).

|  |  |  | CH<sub>4</sub> |  | CO<sub>2</sub> |  | N<sub>2</sub>O |  | CO<sub>2</sub>e |
| --- | --- | --- | --- | --- | --- | --- | --- | --- | --- |
| Article | Group | Time (d)<sup>a</sup> | Flux | Traj. | Flux | Traj. | Flux | Traj. |  |
| Penttilä et al. 2013<sup>§</sup> | DO | 50 | >-100x | +BC | -2.8x | -L | ~1x | +G | Follows CO<sub>2</sub> trajectory |
| Iwasa et al. 2015<sup>¥</sup> | DO | 6 | -2x | -L | -5x | -L | +2x | -BC |  |
| Slade et al. (2016ab)<sup>§</sup> | DC | 60,60 | >-1000x | +BC | -10.67 | +BC | +5x | +G |  |
| Hammer et al. 2016 | DO | 43 | - |  | - |  | - |  |  |
| Piccini et al. 2017<sup>¥</sup> | DC | 32 | >-25x | -L | -3x | -E | >-13x | -E |  |
| Evans et al. 2019<sup>¥</sup> | DO | 56 | ~1x | A | -1.36x | -BC | -2.5x | S |  |
| Fowler et al. 2020c<sup>¥</sup> | DC | 7,14 | >-200x | -L | >-2x | -L | +1.25x | +E |  |

BC, Skewed Bell Curve; L, Linear; G, Gaussian; E, Exponential; S, Sine; A, Alternating.
<sup>a</sup> Calculated difference of groups that represent the ratio of the means of (Day 0)/(DayX), where DayX=Time (d) reported here.
<sup>§</sup> Statistical tests not reported in the published paper thus ‘significant’ differences based on visual means and standard errors as shown in graphs.
<sup>¥</sup> Omnibus ANOVA’s reported ( $p < 0.05$ ).

**Table 4.**
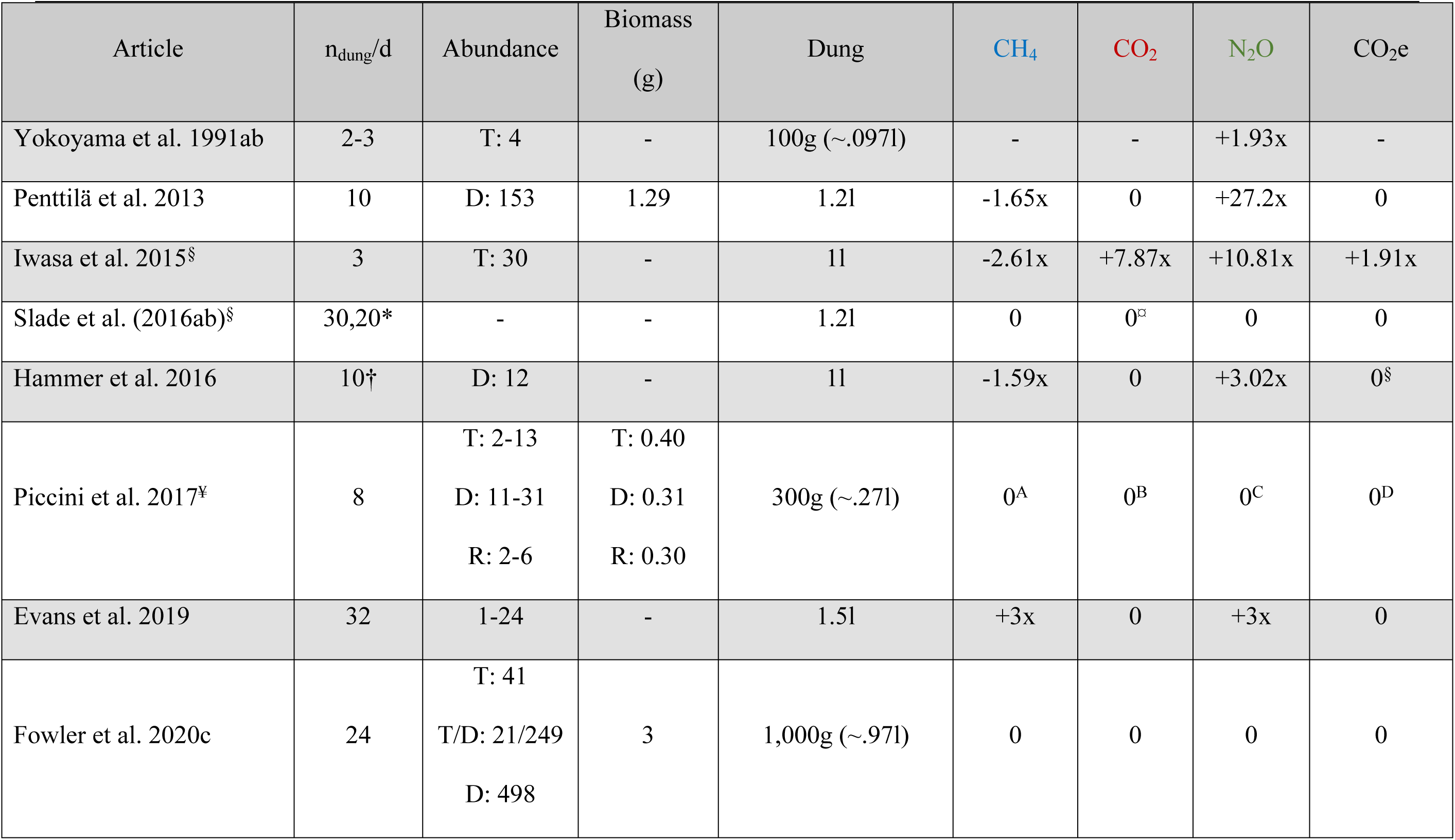
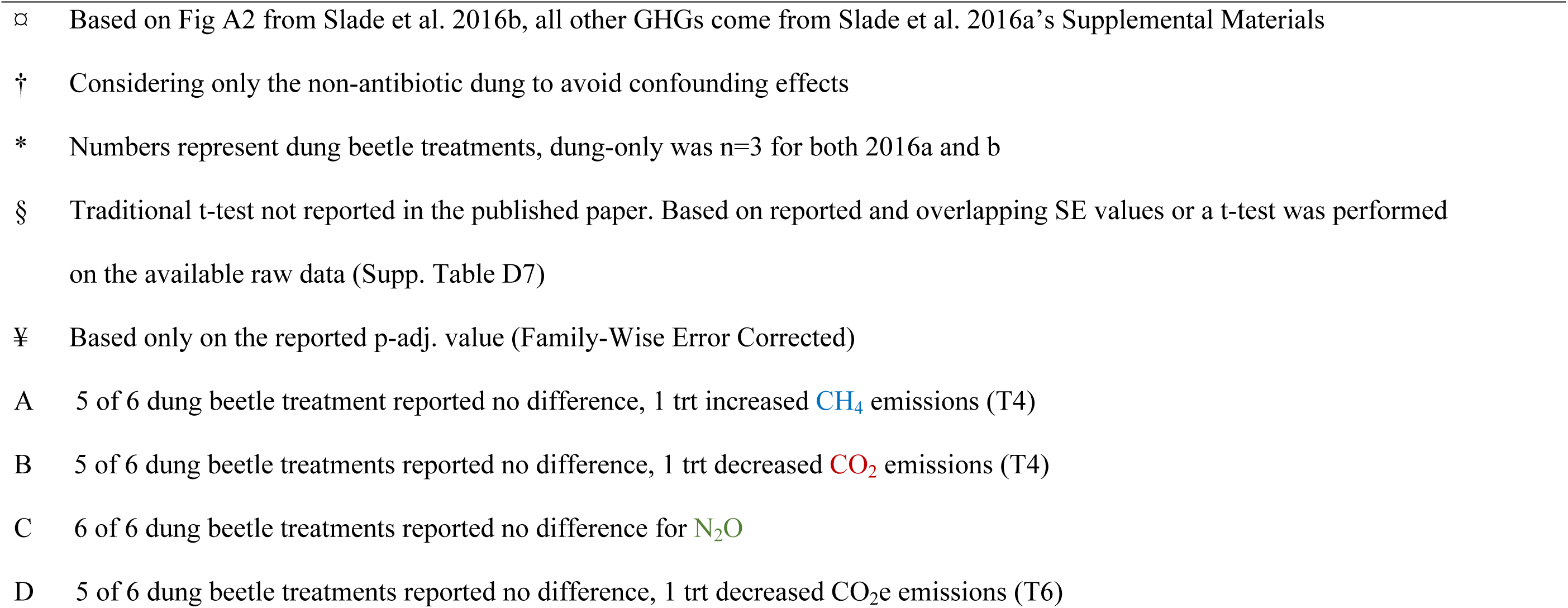
A summary of reported GHG differences (+: increase, -: decrease, 0: no effect) focused on aggregate treatment effects of t-tests and ANOVAs (p<0.05) that exclude strong and effect-masking predictors such as time or vegetation.

| Article | n<sub>dung</sub>/d | Abundance | Biomass<br>(g) | Dung | CH<sub>4</sub> | CO<sub>2</sub> | N<sub>2</sub>O | CO<sub>2</sub>e |
| --- | --- | --- | --- | --- | --- | --- | --- | --- |
| Yokoyama et al. 1991ab | 2-3 | T: 4 | - | 100g (~.097l) | - | - | +1.93x | - |
| Penttilä et al. 2013 | 10 | D: 153 | 1.29 | 1.2l | -1.65x | 0 | +27.2x | 0 |
| Iwasa et al. 2015<sup>§</sup> | 3 | T: 30 | - | 1l | -2.61x | +7.87x | +10.81x | +1.91x |
| Slade et al. (2016ab)<sup>§</sup> | 30,20* | - | - | 1.2l | 0 | 0<sup>□</sup> | 0 | 0 |
| Hammer et al. 2016 | 10<sup>†</sup> | D: 12 | - | 1l | -1.59x | 0 | +3.02x | 0<sup>§</sup> |
| Piccini et al. 2017<sup>¥</sup> | 8 | T: 2-13<br>D: 11-31<br>R: 2-6 | T: 0.40<br>D: 0.31<br>R: 0.30 | 300g (~.27l) | 0<sup>A</sup> | 0<sup>B</sup> | 0<sup>C</sup> | 0<sup>D</sup> |
| Evans et al. 2019 | 32 | 1-24 | - | 1.5l | +3x | 0 | +3x | 0 |
| Fowler et al. 2020c | 24 | T: 41<br>T/D: 21/249<br>D: 498 | 3 | 1,000g (~.97l) | 0 | 0 | 0 | 0 |

- - Based on Fig A2 from Slade et al. 2016b, all other GHGs come from Slade et al. 2016a's Supplemental Materials - † Considering only the non-antibiotic dung to avoid confounding effects - \* Numbers represent dung beetle treatments, dung-only was n=3 for both 2016a and b - § Traditional t-test not reported in the published paper. Based on reported and overlapping SE values or a t-test was performed on the available raw data (Supp. Table D7) - ¥ Based only on the reported p-adj. value (Family-Wise Error Corrected) - A 5 of 6 dung beetle treatment reported no difference, 1 trt increased CH<sub>4</sub> emissions (T4) - B 5 of 6 dung beetle treatments reported no difference, 1 trt decreased CO<sub>2</sub> emissions (T4) - C 6 of 6 dung beetle treatments reported no difference for N<sub>2</sub>O - D 5 of 6 dung beetle treatments reported no difference, 1 trt decreased CO<sub>2</sub>e emissions (T6) -

In our study, CH_4_, CO_2_, and CO_2_e explained 84%, 72%, and 79% of the total variation (8-10x the variation explained by treatment alone). Only for N_2_O did the time:treatment interaction fit 1.64x more variation than either variable alone, explaining a total of 17% variation. This suggests that time, rather than treatment, affects the interaction effect most strongly (except in the case of N_2_O) which explains why dung beetles in our study affected only one out of five sampling days (Fig 5) – a miniscule effect. It is likely that time may possess such a strong effect because it is multicollinear: other variables such as dung moisture, soil moisture, decay, and other unknown time-related variables drastically and simultaneously change (Supp. Table D8) as the pat ages (Fig 2). However, by comparing dung beetle treatments to time, we can more easily deduce if dung beetle activity, despite all other pressures, is a more powerful GHG predictor – it wasn’t. We also hypothesized that premixing dung, an activity reflecting dung beetle activity, might obscure the dung beetle effect – it didn’t.

### Mixing the Message

While the unmixed dung-only saw minor increases in CH_4_, N_2_O, and CO_2_e relative to the mixed dung-only (Figs 5 and 6), these differences did not affect aggregate fluxes (Fig 3) nor did they compare to time’s strong effects (Fig 4). In fact, Evans et al. (2019)’s suggestion that dung tunnels may increase CH_4_’s release from the pat applies only to dung wet enough for anaerobic maintenance, but chunky/dry enough to support sturdy microtunnels. When mixing fresh dung, the dung reforms and reconnects when wet, so likely there was no aeration in the mixed dung-only without O_2_ for confirmation. At the outset we hypothesized that mixing multiple dung pats and relocating them alters GHGs, but this was not borne out – however, it does question whether mixing itself (Table 4) is an influential factor, especially compared to time-based decay (Table 3).

Thus, studying dung beetle populations that can accelerate dung decay faster than time alone may answer what behavior(s) strongly alter GHG pathways. Varying dung beetle abundances in future studies may answer this question.

## Conclusion

Our major findings revealed that: 1) dung-presence always increased GHG (CH_4_, CO_2_, N_2_O) production relative to vegetation-only – likely because the sudden deposition of a rich and readily available nitrogen, carbon, and mineral source (fertilization) sparked microbial activity in the form of gases; 2) that time was the single strongest predictor of GHG trends for reducing CH_4_, CO_2_, and CO_2_e, and steadily increasing N_2_O over time with the potential help of burial activity. These trends may generally be explained by decreased dung moisture, increased soil moisture, and/or dung beetle tunneling behavior that exposes dung to greater oxygenic/microbial consumptive conditions; and 3) that neither physically mixing nor dung beetle activity, when compared with time or in aggregate, affected the total greenhouse gas effect (CO_2_e) in a practical manner, though dung beetles periodically decreased CH_4_ and increased N_2_O. Thus, while dung beetles occasionally influenced individual GHGs in small ways (Table 4), dung-presence and time was a much stronger predictor (Table 3), and thus forces researchers to ask: what kind of impact would we want to see from dung beetles? Future research may help answer whether dung beetles have no GHG effect or if an effect is only observed at greater abundances and activities than currently seen.

## Acknowledgements

Thank you, Amelia Weaver and Tashiana Wilcox for assisting with field experiments and beetle capture. We appreciate Cong Tu for conveying the basic concepts of GHG collection and measurement, as well as the NCSU Lake Wheeler Road Field Lab for collaborating with us on cattle pasture use. Lastly, we appreciate the crucial edits and thoughts of Meredith Spence Beaulieu and Bradley Mullens in the drafting of this manuscript.

This material is based upon work supported by the National Science Foundation Graduate Research Fellowship Program under Grant No. (DGE-1746939). Any opinions, findings, and conclusions or recommendations expressed in this material are those of the author(s) and do not necessarily reflect the views of the National Science Foundation. This research was also supported by the Center for Environmental Farming Systems Graduate Student Fellowship at North Carolina State University.

S1 File. Supplementary Materials.

